## Supplemental Figures for "Ordered dephosphorylation initiated by the selective proteolysis of cyclin B drives mitotic exit"

### SUPPLEMENTAL FIGURE LEGENDS

**Figure 2-figure supplement 1. An ordered cascade of dephosphorylation in anaphase A and B.**

HeLa cells were arrested in mitosis and mitotic exit triggered by MPS1 inhibition with AZ3146 (Hayward et al., 2019). Heatmap of (A) protein groups or (B) phospho-sites clustered based on trends over time (Figure 2-source data 3). Time is shown in minutes. (C) Sequence logo analysis showing the relative enrichment of residues in the adjacent cluster from (B). The number of phospho-sites and average half-life of each cluster is indicated. (D) Key linking heatmap colour with observed ratio. Quantitative ratio of (E-F) CCNB1 and PRC1 protein level or (G) PPP1CA-pT320 and PRC1-pT481 phosphorylation (Mean±SEM, n=2).

**Figure 3-figure supplement 1. Clustering analysis of fractionated anaphase proteomes.**

(A) Heatmap of clustered protein groups generated from fractionated total proteome samples (n=2) (Figure 3-source data 1). (B) Key linking heatmap colour with observed ratio. Quantitative ratio of the protein levels of (C) CCNB1 (Mean±SEM, n=2) , (D) CDC20 (n=1) and (E) Geminin (GMNN) (n=1).

**Figure 4-figure supplement 1. Ongoing protein synthesis is not required to maintain spindle assembly checkpoint-dependent arrest during mitosis.**

(A) Live cell imaging of endogenously CRISPR-tagged CCNB1, following addition of 100 ng/ml nocodazole with either DMSO or 100 µg/ml cycloheximide (CHX). (B) Raw fluorescence intensity plot of cells over time, following drug addition (DMSO n=12, CHX n=6). Only cells which enter mitosis are shown on the graph. Images shown are sum intensity projections, quantification was carried out on sum intensity projections. All times shown are relative to the point of drug addition.(C-D) Cells were arrested in mitosis using (C) nocodazole or (D) taxol, prior to the addition of either DMSO or 100 μg/ml CHX, to inhibit protein synthesis. Changes in protein level were then followed by Western blot. Densitometric quantification of Western blot data from (C-D) for (E) CDC20 or (F) CCNB1 (Mean±SEM, n=3). (G) Statistical analysis of the time taken for cells from Figure 4A to initiate cleavage furrow ingression following chromosome separation using a Mann-Whitney test, * denotes the calculated p-value.

**Figure 6-figure supplement 1. High-resolution analysis of mitotic exit following APC/C inhibition.**

Western blot of samples generated from mitotically arrested cells HeLa following treatment with either (A) DMSO or (B) APC/C inhibitors (APC/Ci), 400 μM apcin and 25 μM pro-TAME, prior to CDK inhibition (CDKi). Samples were taken at the indicated timepoints.

### FIGURE SUPPLEMENTS

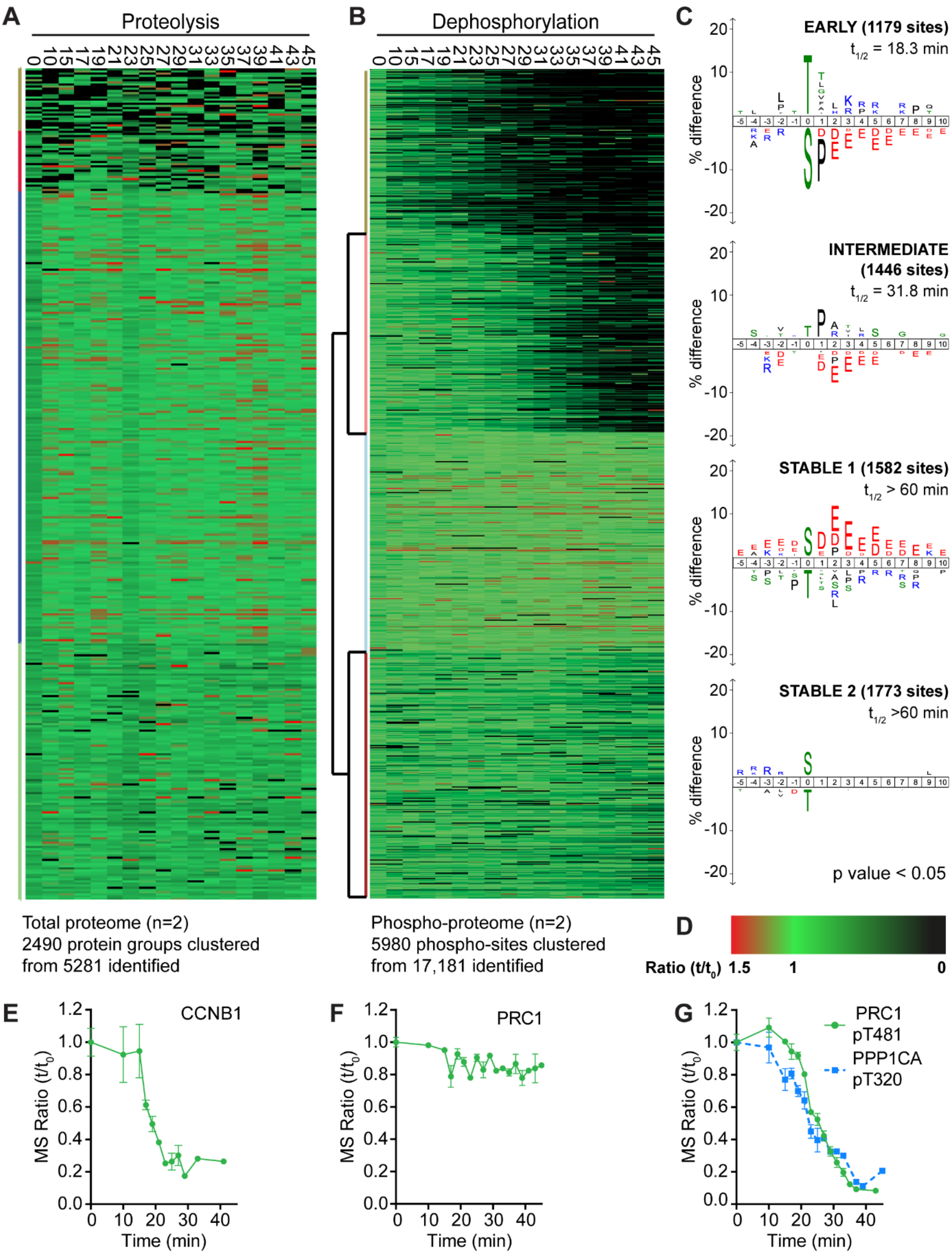


**Figure 2-figure supplement 1. An ordered cascade of dephosphorylation in anaphase A and B.**

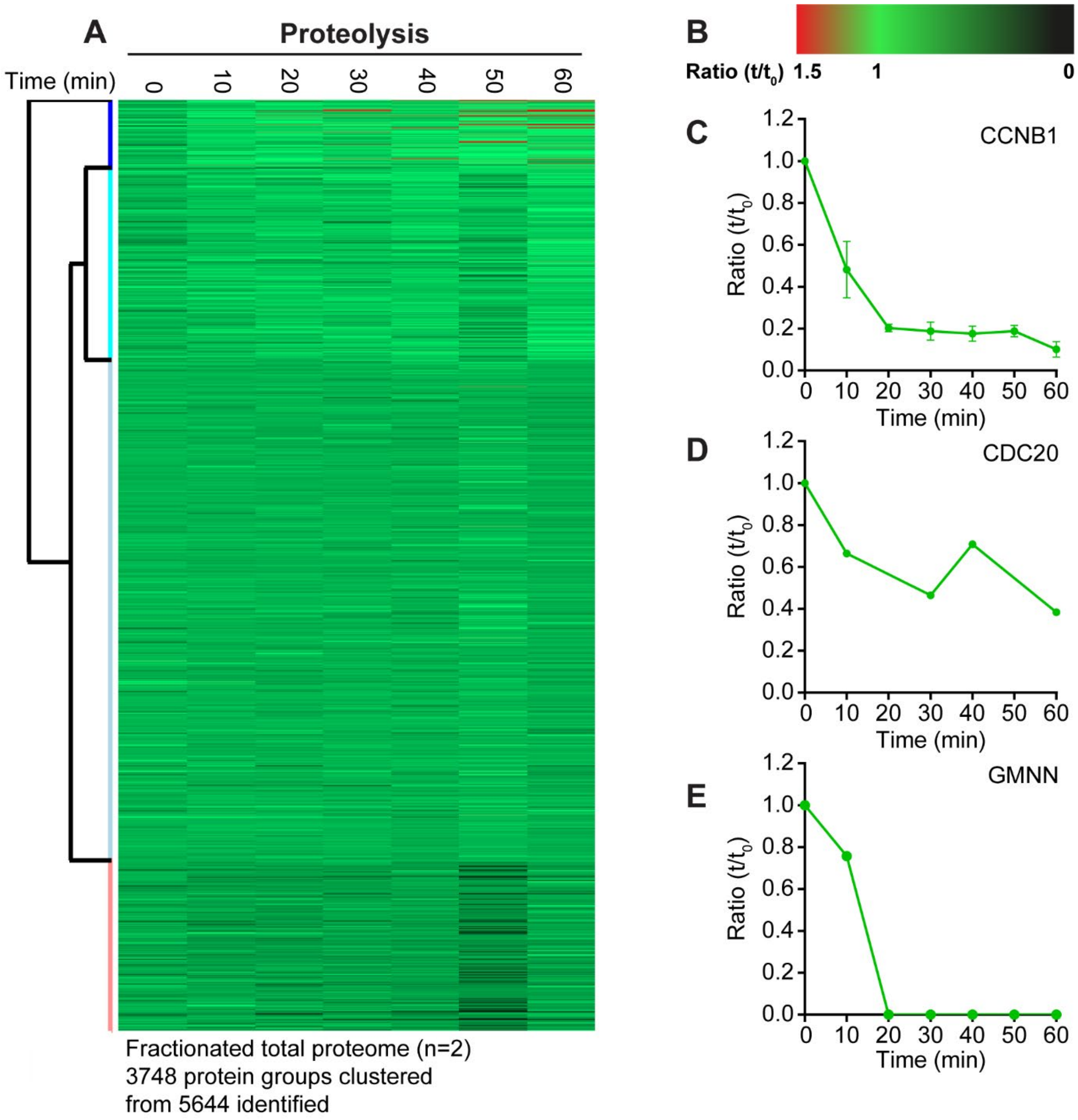


**Figure 3-figure supplement 1. Clustering analysis of fractionated anaphase proteomes.**

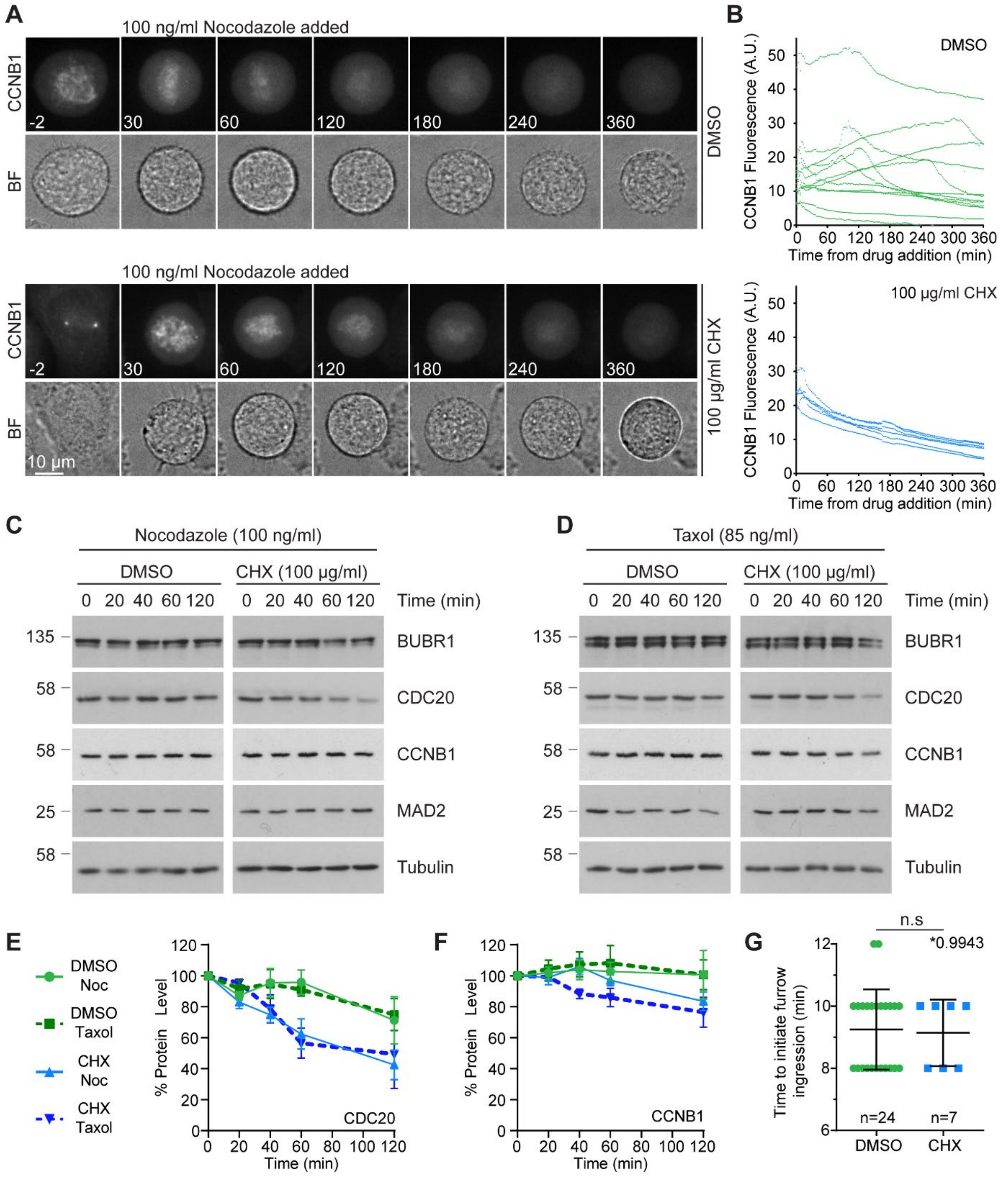


**Figure 4-figure supplement 1. Ongoing protein synthesis is not required to maintain spindle assembly checkpoint-dependent arrest during mitosis.**

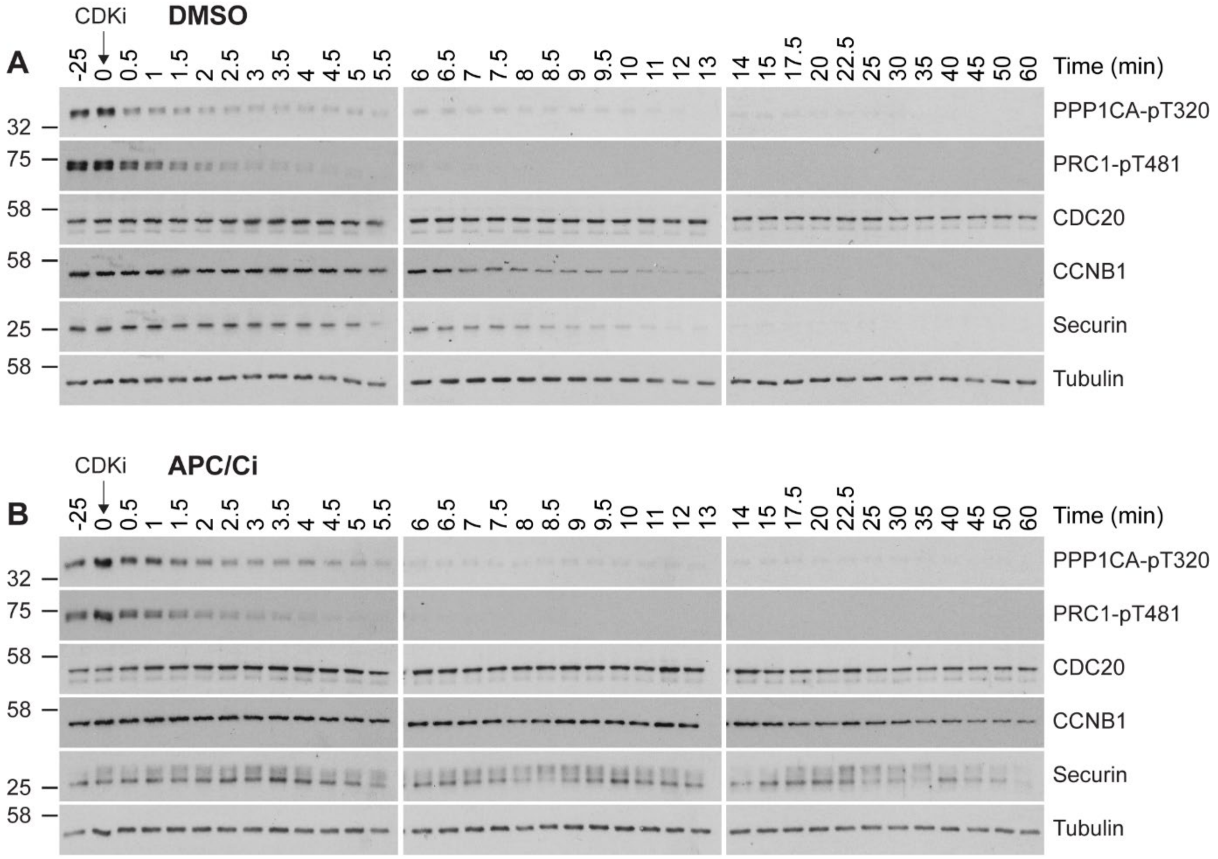


**Figure 6-figure supplement 1. High-resolution analysis of mitotic exit following APC/C inhibition.**
